# Characterizing the floral resources of a North American metropolis using a honey bee foraging assay

**DOI:** 10.1101/834804

**Authors:** Douglas B. Sponsler, Don Shump, Rodney T. Richardson, Christina M. Grozinger

## Abstract

Roughly a third of described insect species visit flowers, making the flower-insect interface one of the chief pillars of global biodiversity. Studying flower-insect relationships at the scale of communities and landscapes has been hindered, however, by the methodological challenges of quantifying landscape-scale floral resources. This challenge is especially acute in urban landscapes, where traditional floral surveying techniques are ill-suited to the unique constraints of built environments. To surmount these challenges, we devised a “honey bee foraging assay” approach to floral resource surveying, wherein continuous colony weight tracking and DNA metabarcoding of pollen samples are used to capture both the overall availability and taxonomic composition of floral resources. We deploy this methodology in the complex urban ecosystem of Philadelphia, PA, U.S. Our results reveal distinct seasonality of floral resource availability, with pulses of high availability in May, June, and September, and a period of prolonged scarcity in August. Pollen genus richness mirrored this pattern, with peak richness in May and June. The taxonomic composition of pollen samples varied seasonally, reflecting underlying floral phenology, with especially strong turnover between May and June samples and between August and September samples delineating well-defined spring, summer, and fall floral resource communities. Trait analysis also revealed marked seasonal structure, with spring samples characterized by trees and shrubs, summer samples including a stronger presence of herbaceous “weeds”, and fall samples dominated by woody vines. Native flora predominated in spring, giving way to a preponderance of exotic flora in summer and fall. Our study provides a detailed portrait of floral resources in a complex urban environment. At a basic level, this yields insight into the assembly of novel urban floral resource communities, showcasing, for example, the emergence of a woody-vine-dominated fall flora. At an applied level, our data can inform urban land management, such as the design of ecologically functional ornamental plantings, while also providing practical guidance to beekeepers seeking to adapt their management activities to floral resource seasonality. Methodologically, our study demonstrates the potential of the honey bee foraging assay as an efficient and standardizable technique for landscape-scale floral resource surveying.

## Introduction

It has been estimated that roughly a third of all described insect species are either directly or indirectly dependent on flowers for food (Wardhaugh 2015). This ecological centrality of flowers-as-food extends to systems in which historic floral communities have been dramatically altered, such as urban landscapes characterized by novel assemblies of native and exotic flora (Aronson et al. 2014). Urban landscapes can host diverse communities of flowers and flower-visitors, sometimes even functioning as refugia for rare species (Baldock et al. 2015) [though see Geslin et al. (2013)], but little is known about the overall composition of urban floral resources and how they fluctuate phenologically through the growing season.

Studying floral resources at the landscape scale is technically daunting in any context (Frankl et al. 2005), but the challenge becomes especially acute in urban landscapes where extreme landscape heterogeneity, limited land access, and the physical obstacles of the built environment render traditional approaches to floral surveying impracticable. Flower-visiting insects might be harnessed as efficient environmental samplers of landscape-scale floral resources, provided the spatial and taxonomic scope of their interaction with the landscape is sufficiently understood and the materials they collect can be analyzed informatively. While not an unbiased representation of local flora, such an approach would manifestly be relevant to the question of trophic function at the flower-insect interface.

The western honey bee (*Apis mellifera* L.) is arguably the organism best suited for such a sampling approach. As an extreme generalist, honey bees have a diet breadth that overlaps considerably with that of many other nectar- and pollen-feeding insects (Goulson 2003). A honey bee colony surveys the landscape around its nest at a range routinely extending several kilometers, dynamically allocating foraging effort to the most rewarding patches (Visscher and Seeley 1982). Thus, the materials it collects are informatively biased toward the richest resources within the colony’s foraging range and compatible with honey bee behavior and morphology (excluding, for example, flowers requiring sonication to release pollen from anthers).

Here, we present a two-year study of the floral resource composition and dynamics of Philadelphia, Pennsylvania, U.S. Founded in 1682, the city of Philadelphia straddles the interface of the Appalachian Piedmont and Atlantic Coastal Plain physiographic provinces (Fenneman and Johnson 1946) and hosts over 2,500 plant species (Clemants and Moore 2003). Its patchwork heterogeneity of sociological composition is both a product and driver of concomitant ecological variation in abiotic substrate and biotic communities, earning Philadelphia the apt moniker “City of Neighborhoods”. To characterize the floral resources of this complex urban environment, we employ a “honey bee foraging assay” using a network of sentinel apiaries distributed throughout the city. Combining DNA metabarcoding of pollen samples with continuous colony weight monitoring, we characterize both the taxonomic composition and overall availability of floral resources in our study system, together with their temporal dynamics throughout the foraging season. We conclude by comparing Philadelphia’s floral resource landscape with patterns described in other study systems and discussing the implications of our data for understanding the ecological function of urban flora.

## Methods

### Sites and study years

We conducted field work in 2017 and 2018. Our study sites included 13 apiaries in Philadelphia, PA, USA, owned and managed by the Philadelphia Bee Company **(Figure 1)**. Twelve apiaries were used in each study year, with 11 shared across years, 1 used only in 2017 (RB), and 1 used only in 2018 (NE). Each of our research apiaries included three honey bee colonies designated as research colonies for our study, resulting in a total of 36 research colonies across 12 sites in each year. Some sites also included additional colonies not involved in our study. Research colonies were initiated in April-May of each study year from 4- or 5-frame nucleus colonies installed in 10-frame Langstroth hives. During colony installation, each research hive was equipped with a Sundance I bottom-mounted pollen trap (Ross Rounds,) and a Broodminder hive scale (Broodminder, Stoughton, WI).

**Figure 1:**
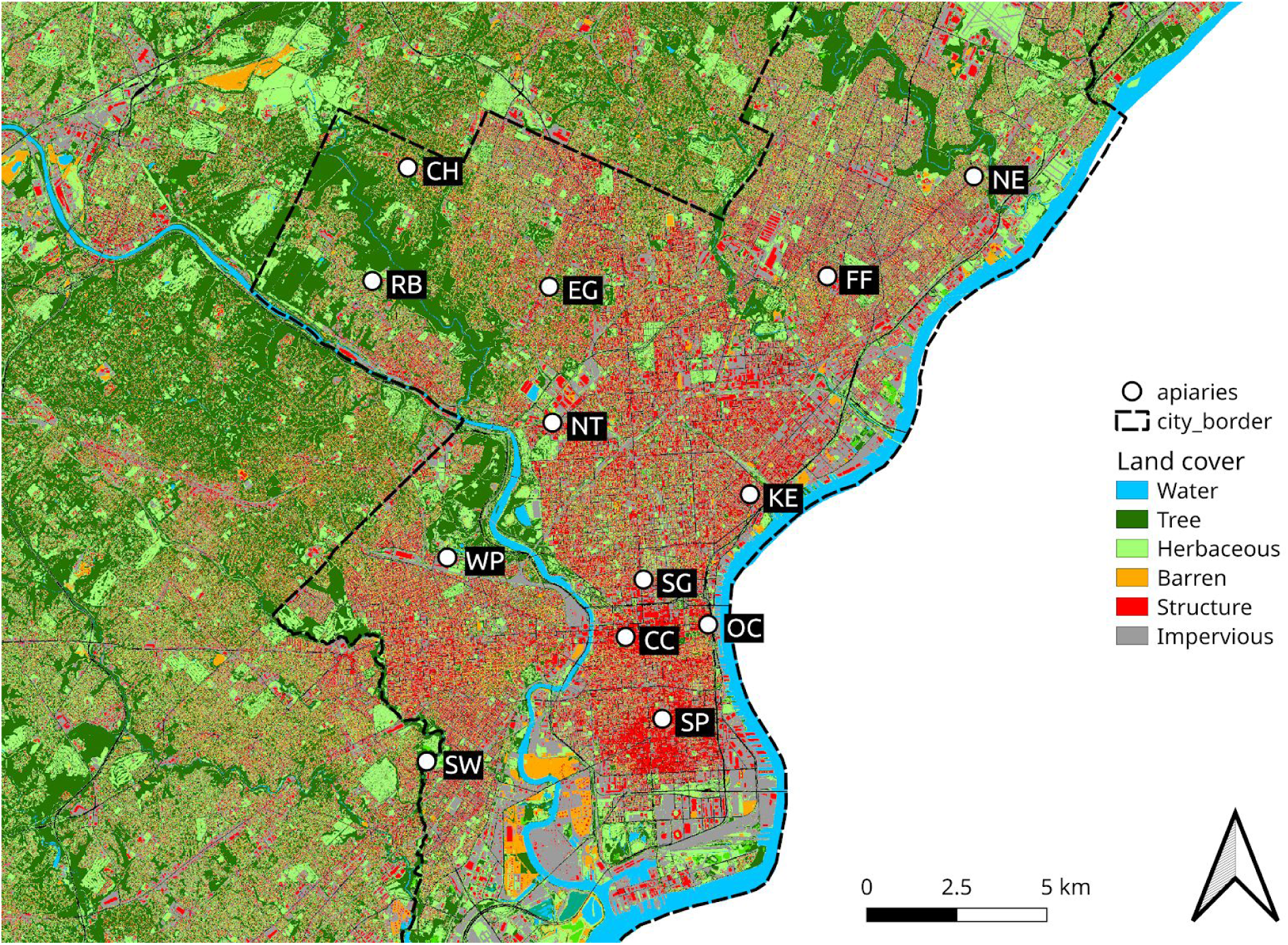
Apiary locations (circles) plotted over land cover raster. The City of Philadelphia is demarcated by a dashed black line. Land cover data are from the Chesapeake Conservancy’s Land Cover Data Project (Chesapeake Bay Conservancy 2017); only major cover types are shown in legend. Area east of the Delaware River in the State of New Jersey is not shown because it fell outside the scope of the land cover data set.

### Weight monitoring and pattern characterization

We set our Broodminder hive scales to record hourly weights, beginning at colony installation in the spring of each year and continuing through the end of our sampling season in the fall of each year. During preliminary data visualization, we identified two types of artifacts in our weight data: apparently random misreadings, resulting in transient (single reading) weight spikes (i.e. additive outliers), and persistent weight shifts (i.e. level-shift outliers) caused by colony manipulations involving the addition or removal of material from the hive. Because our colonies were part of a working apicultural business, it was not possible to avoid or standardize these management artifacts.

To address both types of artifacts, we first took the first-order difference of each colony weight time series, which turns level-shift outliers into additive outliers. We then applied a filter to each time series where data points (i.e. weight changes between consecutive readings) with an absolute value of 2.5 kg or more where identified as outliers and set to zero. The threshold of 2.5 kg was chosen because histograms of first-order differenced time series showed consistent discontinuities around +/-2.5 kg, such that the interval −2.5 to 2.5 contained a smooth curve of ostensibly “normal” readings, while readings outside this interval were rare and extreme. We then reintegrated each time series to reconstruct de-artifacted weight curves and selected only midnight readings for downstream analysis. For the purposes of our study, we were interested principally in the temporal patterns of resource availability arising from landscape-scale floral phenology, and how these patterns might vary site-to-site. To isolate these signals from the noise of colony-level differences in amplitude of weight variation (an expected result of differences in colony strength), we normalized (scaled and centered) the de-artifacted weight curves for each colony. This pipeline will be available upon publication and is depicted graphically in **Figure 2**. Raw data will be available in Appendices S1 and S2 upon publication.

**Figure 2:**
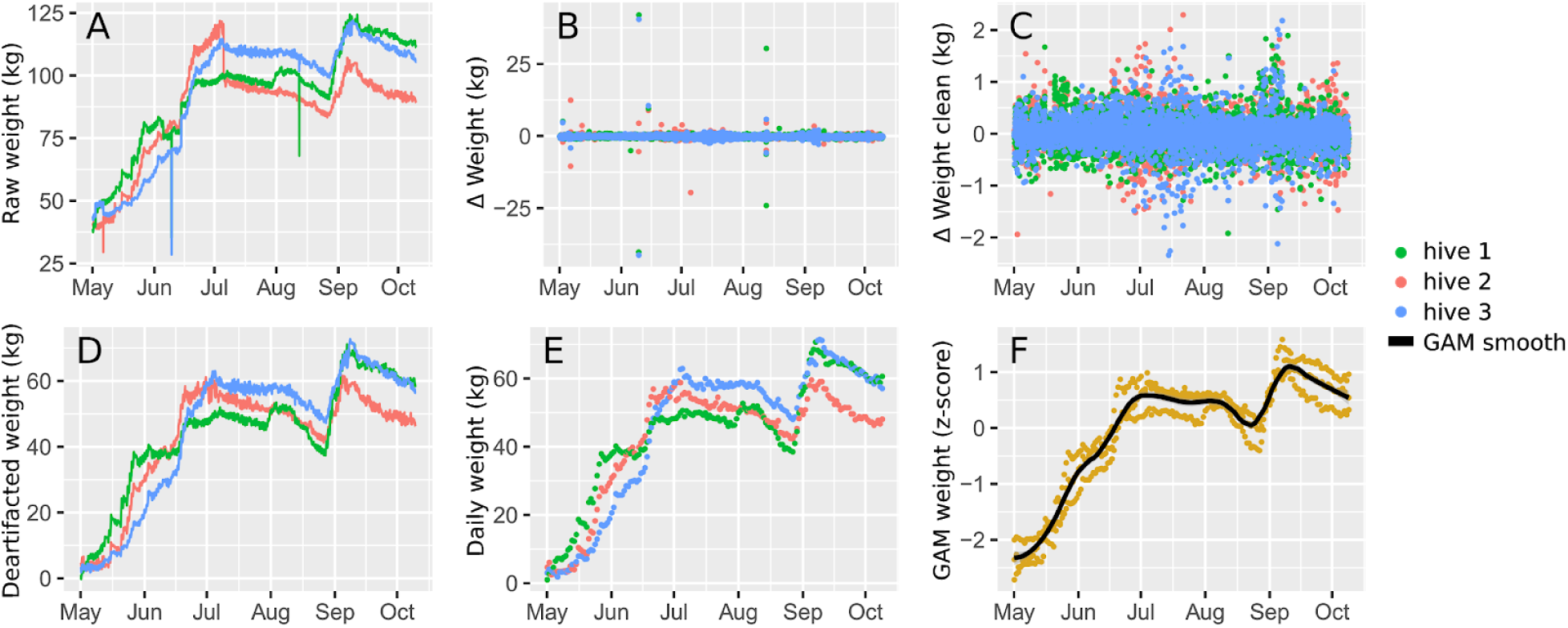
Weight data processing workflow illustrated for three hives at one site. Raw weights (A) contain additive and level-shift outliers. To remove these artifacts, time series were first differenced (B), then filtered (C) and re-integrated (D). Reintegrated time series were them subsampled to include only midnight readings (E). Finally, time series were z-normalized and fit with a GAM smooth (F).

Hives subject to known anomalies were omitted from downstream analysis. In 2017, we omitted all hives from sites CH, EG, and SP due to late colony installation. We also omitted CC hive 3 and RB hive 2 due to scale malfunctioning. In 2018, we omitted SG hive 3 and WP hive 3 due to user error (failure to replace dead batteries), all NE hives due to premature death, and SP hive 3 due to premature death. This left a total of 25 colonies across 9 sites for 2017 and 30 colonies across 11 sites for 2018.

To characterize weight patterns, we used generalized additive modeling (GAM) implemented in R (R Core Team 2019) using package `mgcv` (Wood 2017) to relate colony weight to time and site. Our goals were (1) to identify major gain and loss motifs, (2) to determine whether there was significant site-to-site variation in weight pattern, and (3) to determine whether there was sufficient similarity across sites to justify a global weight model. Accordingly, we constructed three nested hierarchical GAMs, following Pedersen et al. (2019). Model 1 included only a global smooth of weight by time, without site specific smooths. Model 2 included both a global smooth and site-specific smooths. Model 3 included only site-specific smooths, without a global smooth. These models are analogous to the G, GI, and I models, respectively, of Pederson et al. (2019). Since any effect of site or hive on group means was removed by our normalization step, we did not include site or hive random effects. After optimizing the fit of each model, we used AIC model selection to identify the most appropriate model. GAM smooths were then first-order differenced to enable the visualization of detrended gain and loss periods. R code for our GAM analysis will be available in Appendix S8 upon publication.

### Pollen sampling

Pollen samples were collected at roughly monthly intervals. In 2017, sampling occurred in late-May/early-June (hereafter “June sampling period”), late-June/early-July (hereafter “July sampling period”), early/mid-August, and mid-September, for a total of 4 sampling periods. In 2018, we expanded our sampling to include early-May, early-June, early-July, early-August, early-September, and early-October, for a total of 6 sampling intervals. Pollen traps were activated for roughly 3-7 days at the end of each monthly period. Exact sampling dates and durations varied from site to site due to travel time and weather constraints, and a detailed sampling schedule will be available in Appendix S3 upon publication. For the purposes of downstream analysis, sampling dates were binned discretely by monthly sampling period. Samples were collected by scooping the contents of the pollen traps into 50 mL conical tubes. Excess pollen was removed from the trap and poured into the inside of the top box of the hive to help the colonies recover from any pollen deficit caused by our sampling and to ensure that the pollen trap would be empty in preparation for the next sampling period.

### Pollen sample preparation

Pollen processing began with the creation of pooled subsamples by combining 4/*n* g of pollen from each hive within each site-date, where *n* represents the number of hives sampled in a site-date. For most samples, *n* was equal to 3 hives, though there were some cases where fewer than 3 hives at a given site could be sampled, either due to colony death or pollen trap malfunctioning. Each 4 g subsample (hereafter “sample”) was then suspended in 40 mL of 70% EtOH and vortexed until fully dispersed. The resulting suspension was then centrifuged (Eppendorf 5810 R, Hamburg, Germany) at 3000G for 2 minutes and decanted. We then resuspended the pollen sediment in 10 mL of 100% EtOH and repeated the vortexing and centrifugation steps. Lastly, the sample was air dried in an open conical tube under a fume hood for 24 hours to allow the EtOH to evaporate, yielding the final granularized pollen sample.

From each granularized sample, we transferred a 10 mg aliquot to a 2 mL beadmill tube. To each tube, we added 1 mL of either Qiagen buffer AP1 (for 2017 samples) or ultra-pure water (for 2018 samples) and approximately 0.5 mL of 1.3 mm chrome steel beads (BioSpec, BioSpec, Bartlesville, OK). Samples were then subjected to 4 × 1-minute run on an Omni Bead Ruptor 24 Elite (Omni International, Kennesaw, GA) set to a speed of 6 m/s. Samples were iced between runs to prevent overheating.

For our 2017 samples, we purified beadmill lysates using a Qiagen DNeasy Plant Minikit according to the manufacturer’s instructions. In 2018, we streamlined our workflow by using Phire Direct PCR reagents (Thermo Fisher) that allowed us to avoid the purification step (hence the use of water rather than buffer AP1 in beadmilling). In both methods, a 1 μL aliquot of the final product (either purified DNA extract or raw lysate) was used as PCR template.

### Library preparation and sequencing

Past work has shown that using multiple loci in pollen DNA metabarcoding improves the robustness of the method, particularly with respect to quantitative reliability (Richardson et al. 2015, 2019a). For this study, we selected the plastid intron *trnL* and the nuclear ribosomal spacer regions ITS1 and ITS2. Each of these loci has been used individually or in combination with other loci in past applications of pollen DNA metabarcoding (e.g. (Keller et al. 2015, Kraaijeveld et al. 2015, Smart et al. 2017)), though ours is the first study to employ this exact combination. We chose ITS1 and ITS2 because these markers are highly divergent and offer relatively high taxonomic resolution (Chen et al. 2010, Wang et al. 2015); *trnL* is far less divergent but easily amplified (Taberlet et al. 2007), and there is some evidence that single-copy plastid markers might be more quantitatively reliable than the ITS regions (Richardson et al. 2015).

Our PCR methods follow the nested protocol of Richardson et al. (Richardson et al. 2019a), which aims to minimize PCR biases introduced by interactions between template and multiplex indices (Berry et al. 2011). Briefly, PCR1 amplifies barcode target regions (*trnL*, ITS1, and ITS2, in separate reactions), PCR2 adds Illumina read-priming oligonucleotides, and PCR3 adds dual multiplex indices (Kozich et al. 2013, Sickel et al. 2015). For the initial amplification of our target loci (PCR1), we used the trnL-c and trnL-h primers described by Taberlet et al. (2007) and the ITS1-u1, ITS1-u2, ITS2-u3, and ITS2-u4 primers described by Cheng et al. (2016). Following PCR3, we cleaned and normalized our libraries using a SequalPrep normalization kit (Thermo Fisher, Waltham, MA USA), pooling libraries within markers so that they could be sequenced equimolarly. In addition to running negative controls on all gels, we also propagated 3 technical replicate negative controls for each marker through the entire nested PCR protocol and sequenced these alongside our positive samples, enabling us to detect trace contamination that did not appear on gels. We also used a 10% unsaturated matrix of multiplex tags so that critical mistagging rates could be both mitigated and measured (Esling et al. 2015, Schnell et al. 2015). A full description of primers, PCR reaction conditions, and negative controls will be available in Appendices S4, S5, and S6 upon publication.

Prepared libraries were sequenced using 2 × 300 Illumina MiSeq kits, with 2017 and 2018 samples sequenced separately. Sequencing was performed at the Genomics Core Facility of Pennsylvania State University. All sequencing output has been archived on NCBI (available upon publication).

### Bioinformatics

Reference sequences were downloaded from NCBI Genbank on 22 February 2019 using the eDirect API **(Table 1)**. Downloaded reference sequences were then filtered at the genus level to include only genera documented as occurring in the states of Pennsylvania, New York, New Jersey, Delaware, or Maryland based on the USDA PLANTS database (USDA, NRCS 2019). We then used MetaCurator (Richardson et al. 2019b) to trim our reference sequences to the specific regions of our amplicons and taxonomically dereplicate identical sequences.

**Table 1:**
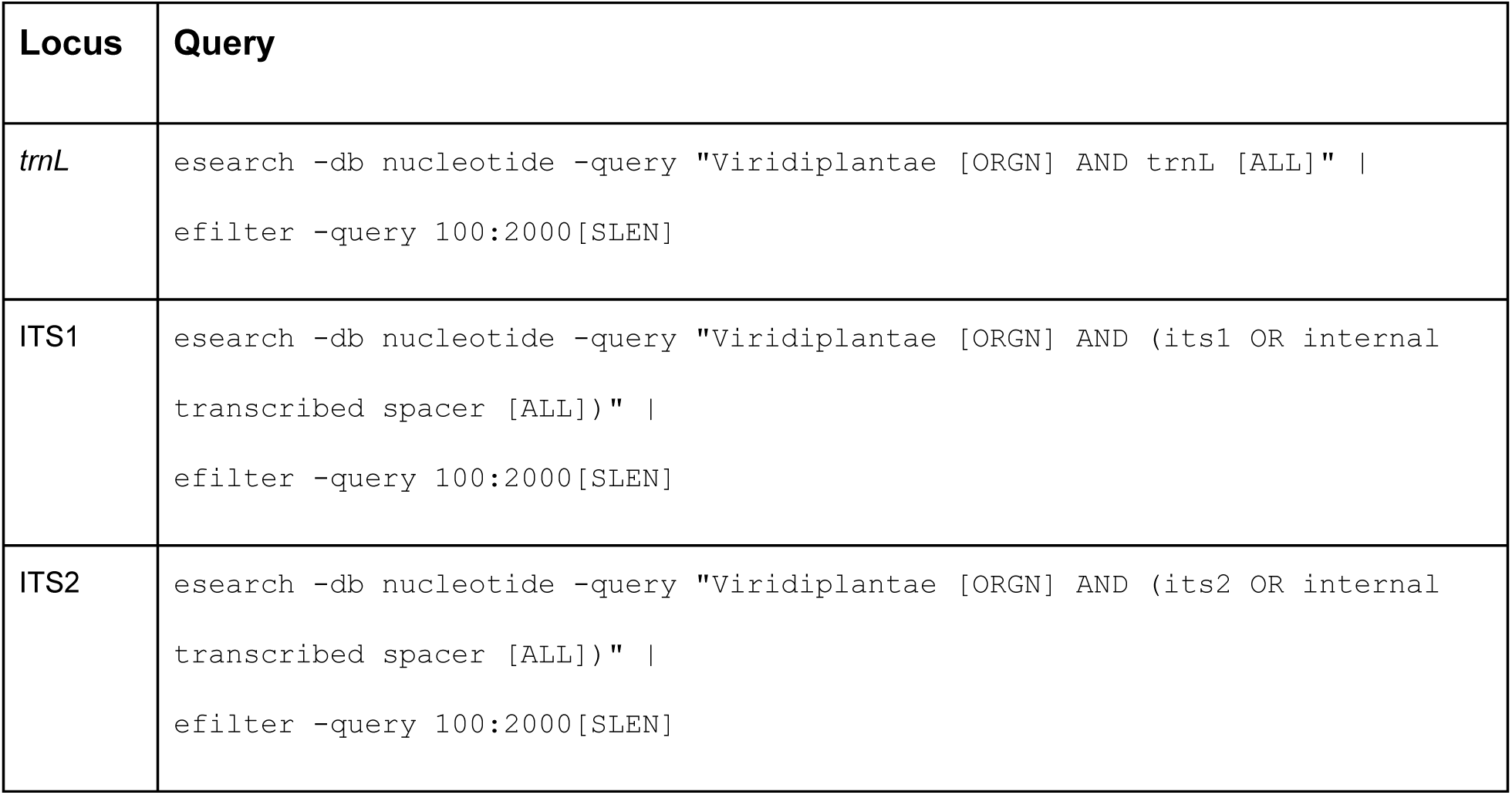
eDirect queries used to download Genbank reference sequences.

MiSeq reads were first mate-paired using PEAR (Zhang et al. 2014) and converted from FASTQ to FASTA format using the FastX Toolkit (Gordon et al. 2010). Mate-paired reads were then aligned to our curated reference sequences using the `usearch_global` algorithm in VSEARCH (Rognes et al. 2016), with a minimum query coverage of 80% and a minimum identity of 75%. Only the top hit for each query was retained for downstream analysis.

After alignments were made, we performed further processing in R (R Core Team 2019). First, alignments were subjected to more stringent identity thresholds: 97% for *trnL*, and 95% for ITS1 and ITS2. Then, subject sequence accession numbers were joined with taxonomic lineages using the R package `taxonomizr` (Scott Sherrill-Mix 2019). Using the grouping and summarizing utilities of the R package `dplyr` (Wickham et al. 2019), we tallied, for each marker, the number of reads by genus and sample (i.e. site-date) and expressed each genus tally as a proportion by dividing its read count by the total number of reads for the sample to which it belonged. Genera comprising less than 0.05% of the total read count for a given sample were discarded to avoid low-abundance false positives. We then used the table joining utilities of `dplyr` (Wickham et al. 2015) to merge the genus tallies across our three markers, and genera detected by only one of the three markers were discarded as potential false positives. Finally, we calculated the median proportional abundance for each genus across all markers by which it was detected and rescaled the median proportional composition of each sample so that it totaled to one. In comparisons with pollen microscopy, this approach has been shown to yield quantitatively informative results (Richardson et al. 2019a). R script for this workflow will be available upon publication.

### Taxonomic analysis

Prior to taxonomic analysis, we excluded any samples in which one of the three libraries were represented by fewer than 1000 reads to avoid making inferences based on insufficient sequencing depth. In 2017, all 44 libraries were adequately sequenced. In 2018, we dropped 6 of 71 libraries (8%) due to under-sequencing, including June samples for sites CH, EG, and NE, September samples for FF and NE, and the October sample for WP.

Using the proportional abundance data described above, we characterized the taxonomic composition of pollen samples across sites, sampling periods, and years using principal coordinates analysis (PCoA) based on the Jaccard dissimilarity metric and implemented with the R package `vegan` (Oksanen et al. 2019). Proportional abundances were square-root transformed prior to ordination, achieving the equivalent of a Hellinger transformation (Legendre and Gallagher 2001) on raw community data. Differences across years, sampling periods, and sites were evaluated by permutation tests using `vegan`’s adonis function (Oksanen et al. 2019). We also analyzed sample genus richness as a function of sampling period. Differences in richness across periods were tested by ANOVA and pairwise differences were analyzed with Tukey’s post-hoc test. R script for this workflow will be available in Appendix S7 upon publication.

### Trait-based analysis

To explore the relationship between sampled relative abundance and genus traits, we classified each genus detected in our samples on the basis of growth habit (tree/shrub, vine, herb), life cycle (perennial, biennial, annual), and native status (native or exotic to contiguous U.S.). Trait data were obtained by downloading from the USDA PLANTS Database (USDA, NRCS 2019) all species records for Philadelphia, PA, along with annotations for “Growth Habit”, “Duration”, and “Native Status”. We opted for a narrower geographical scope for trait analysis than we did for reference sequence curation because the addition of congeners not likely to be present in our local study system would increase intrageneric trait heterogeneity, resulting in more ambiguous trait assignments. Genera found in our pollen data but absent from the USDA PLANTS Database Philadelphia records were manually added to the processed data set by searching the genera in the USDA PLANTS Database without geographical constraints and selecting species with distributions including Pennsylvania, Delaware, New Jersey, Maryland, or New York. Using tidyverse (Wickham 2017) functions in R, we filtered the resulting species list by the genera detected in our pollen samples and summarized each trait for each genus by concatenating unique trait values. Where trait values were not uniform across species within a given genus, the trait was considered “undetermined”. For growth habit, we combined “Tree” and “Shrub” classes into “tree/shrub” and the “Forb/Herb” and “Graminoid” classes in to “herb”. Raw data, derived trait tables, and R script will be available in Appendix S7 upon publication.

## Results

### Weight dynamics

Weight dynamics were characterized by distinct gain and loss motifs that were largely consistent across years, with periods of strong weight gain in May and June, weak gain or weak loss in July, strong loss in August, weak to strong gain in early September, and strong loss from the second week of September to the end of data collection in October **(Figure 3)**. For both years of our study, HGAM analysis with AIC model selection supported the use of both global and site-specific smoothers (Model 2), indicating both a site effect and an archetypal pattern of weight dynamics for our study system **(Table 2)**. Site-specific weight curves including integrated data (i.e. not subject to first-order differencing) will be available in Appendix S8 upon publication.

**Table 2:** AIC output of HGAM analysis.

| Year | Model | Degrees of freedom | AIC |
| --- | --- | --- | --- |
| 2017 | Model 1 (global) | 15 | 6221 |
|  | Model 2 (global + site) | 99 | 5059 |
|  | Model 3 (site) | 105 | 5114 |
| 2018 | Model 1 (global) | 24 | 7132 |
|  | Model 2 (global + site) | 194 | 2923 |
|  | Model 3 (site) | 105 | 5114 |

**Figure 3:**
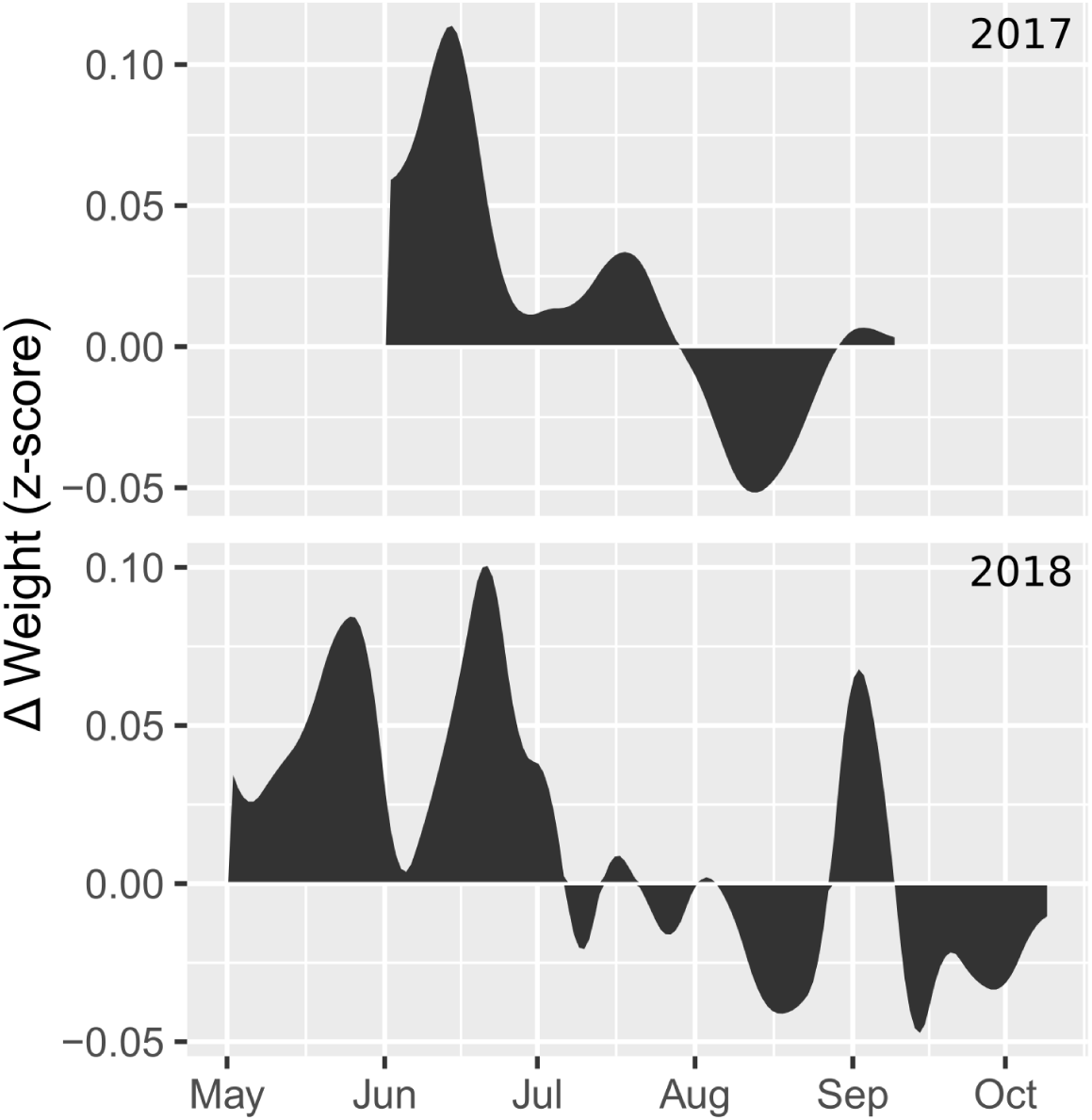
First-order difference of global GAM smooths for 2017 and 2018. Positive and negative y-values reflect weight gain and loss, respectively, in the integrated (non-differenced) data.

### Pollen sample composition

We detected a total of 119 and 139 genera in 2017 and 2018, respectively (tabulated data will be available in Appendices S9 and S10 upon publication). Pollen diversity (genus richness) differed significantly across sampling periods (2017: F = 6.5, p < 0.001; 2018: F = 16.3, p = 0.001), with June having significantly higher diversity (Tukey test, p < 0.05) than August and September in 2017 and May and June having significantly higher diversity (Tukey test, p < 0.05) than all other months in 2018 **(Figure 4)**. A summary of sequencing output by sample will be available in Appendix S11 upon publication.

**Figure 4:**
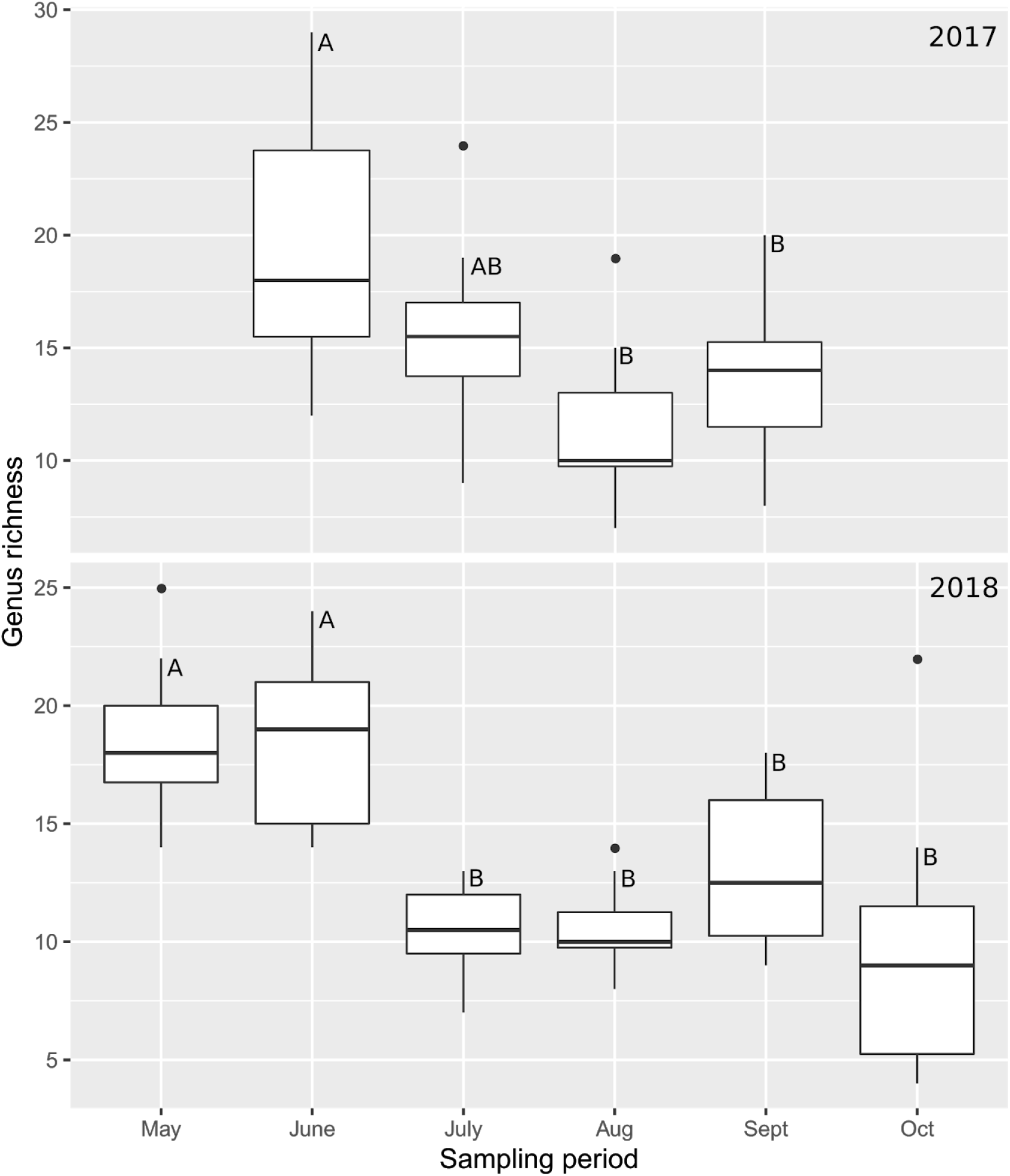
Boxplot of genus richness by sampling period. Letters indicate significant differences according to Tukey’s test.

PCoA revealed strong differentiation across sampling periods in overall pollen composition **(Figure 5)**. The greatest discontinuities were seen across the May-June and August-September intervals. Permutation testing indicated that site (F = 1.5, R^2^ =0.09, p = 0.004), year (F = 5.3, R^2^ =0.03, p = 0.001), and sampling period (F = 16.9, R^2^ =0.43, p = 0.001) were all significant predictors of pollen sample composition, but sampling period explained by far the most variance.

**Figure 5:**
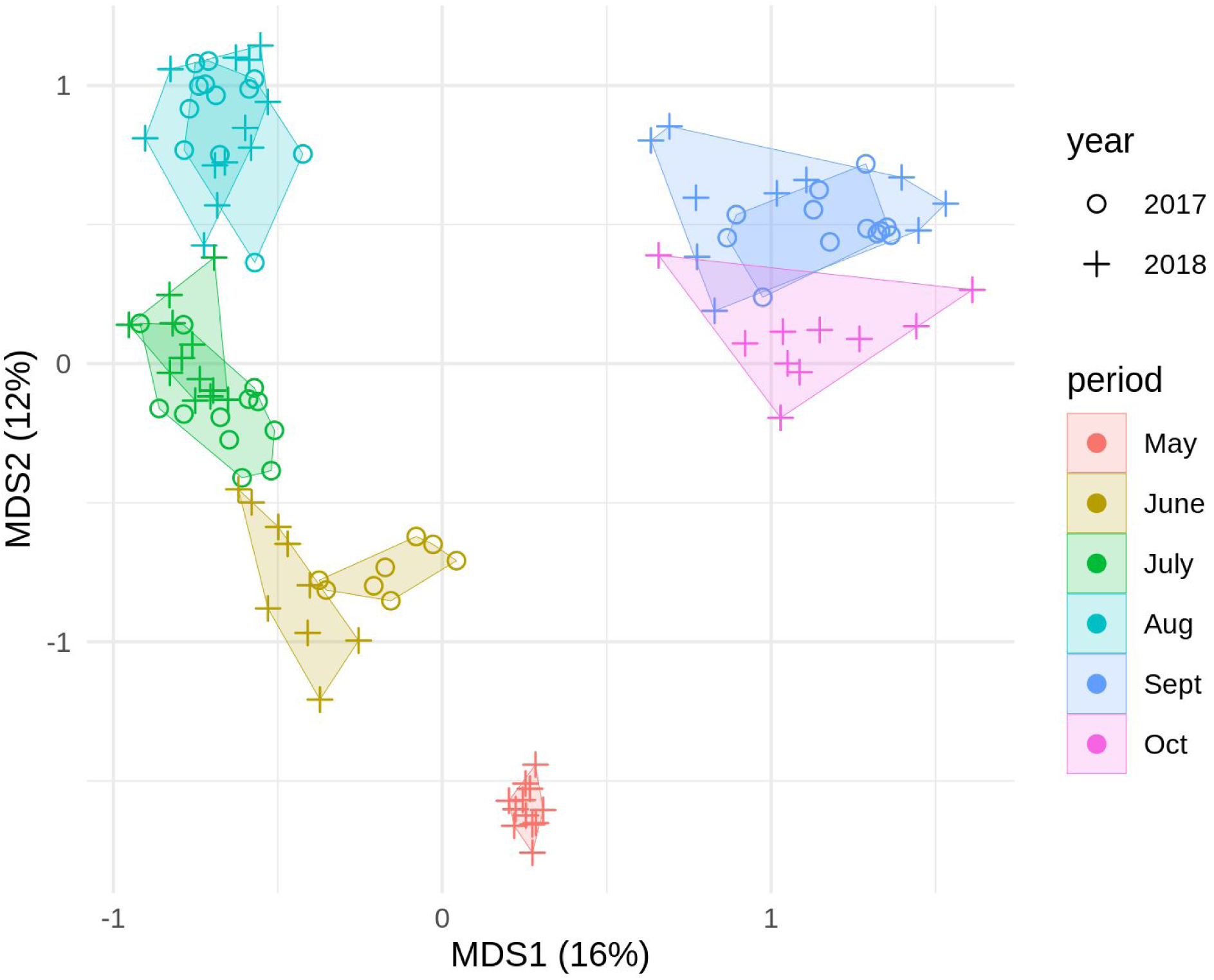
PCoA ordination of pollen data. Pollen samples (site*period) are represented by circles (2017) and crosses (2018). Sampling periods are indicated by color, and samples belonging to each period are joined in convex hulls.

May samples (collected only in 2018) **(Figure 6)** were dominated by wind-pollinated tree genera, including willow (*Salix*), maple (*Acer*), oak (*Quercus*), ash (*Fraxinus*), and plane tree (*Platanus*). Apple (*Malus*) was also abundant. The ornamental shrub Christmas berry (*Photinia villosa* is the only member of its genus present in our study system) dominated the composition of a single site (CC).

**Figure 6:**
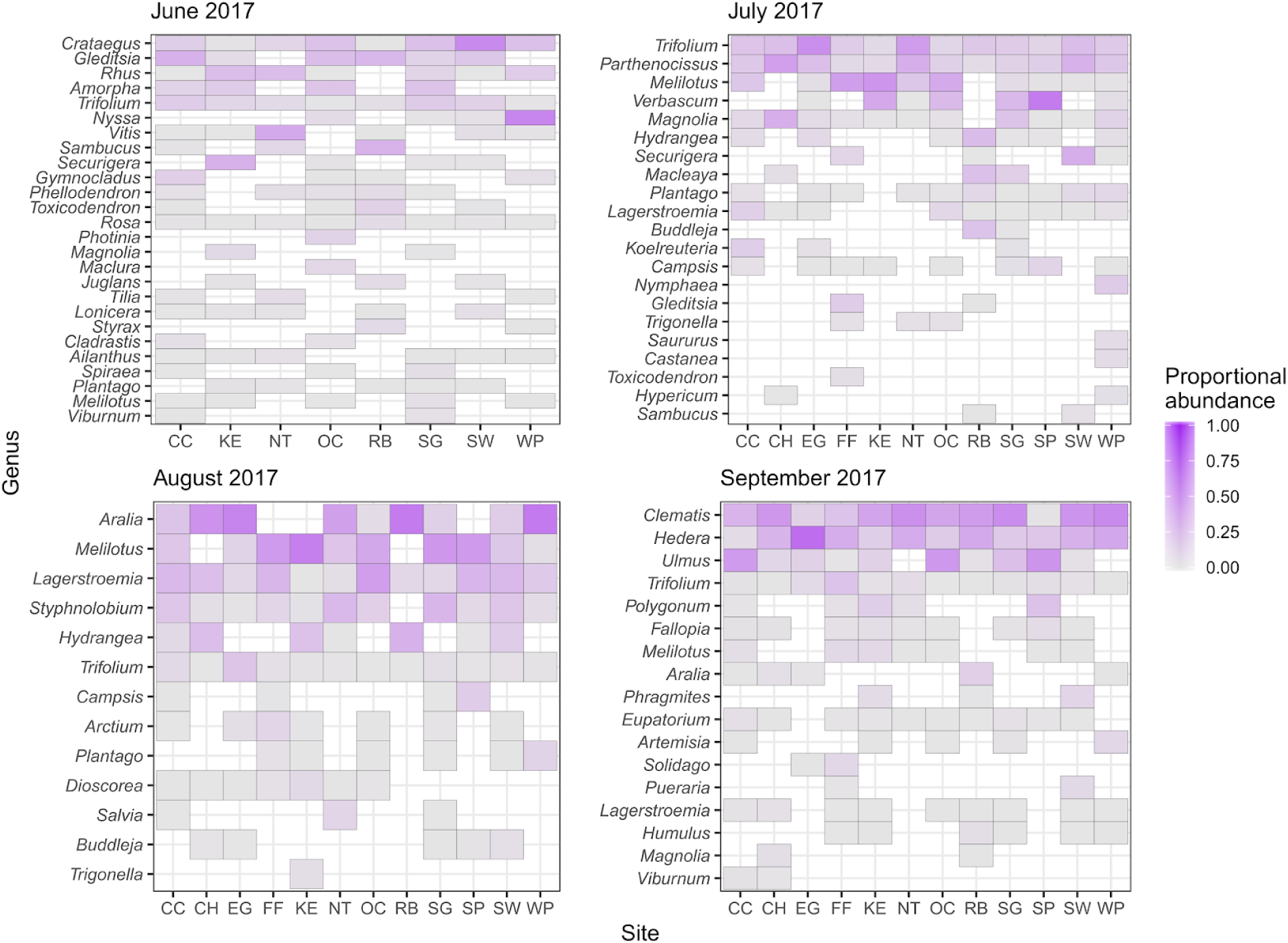
Genus composition of 2017 pollen samples. Genera are ranked by mean proportional abundance across sites. Only genera with a maximum within-site proportional abundance of at least 2.5% are shown.

Between May and June, a strong phenological shift is evident in the floral community, marked by a transient lull in weight gain **(Figure 3)** and a near-total turnover of pollen composition apparent both in the raw data and in the wide discontinuity between May and June samples in PCoA ordination **(Figure 5)**. Following this shift, June pollen samples **(Figures 6, 7)** were characterized by smaller trees and herbaceous flora associated with forest edge, mid-successional, or open landscapes, including hawthorn (*Crataegus*), honey locust (*Gleditsia*), sumac (*Rhus*), clover (*Trifolium*), and sweet clover (*Melilotus*), along with magnolia trees (*Magnolia*) that are common in our study system as ornamental plantings. False indigo (*Amorpha*), a shrub that grows abundantly along the banks of the Schuylkill River, was also abundant in some samples in 2017.

**Figure 7:**
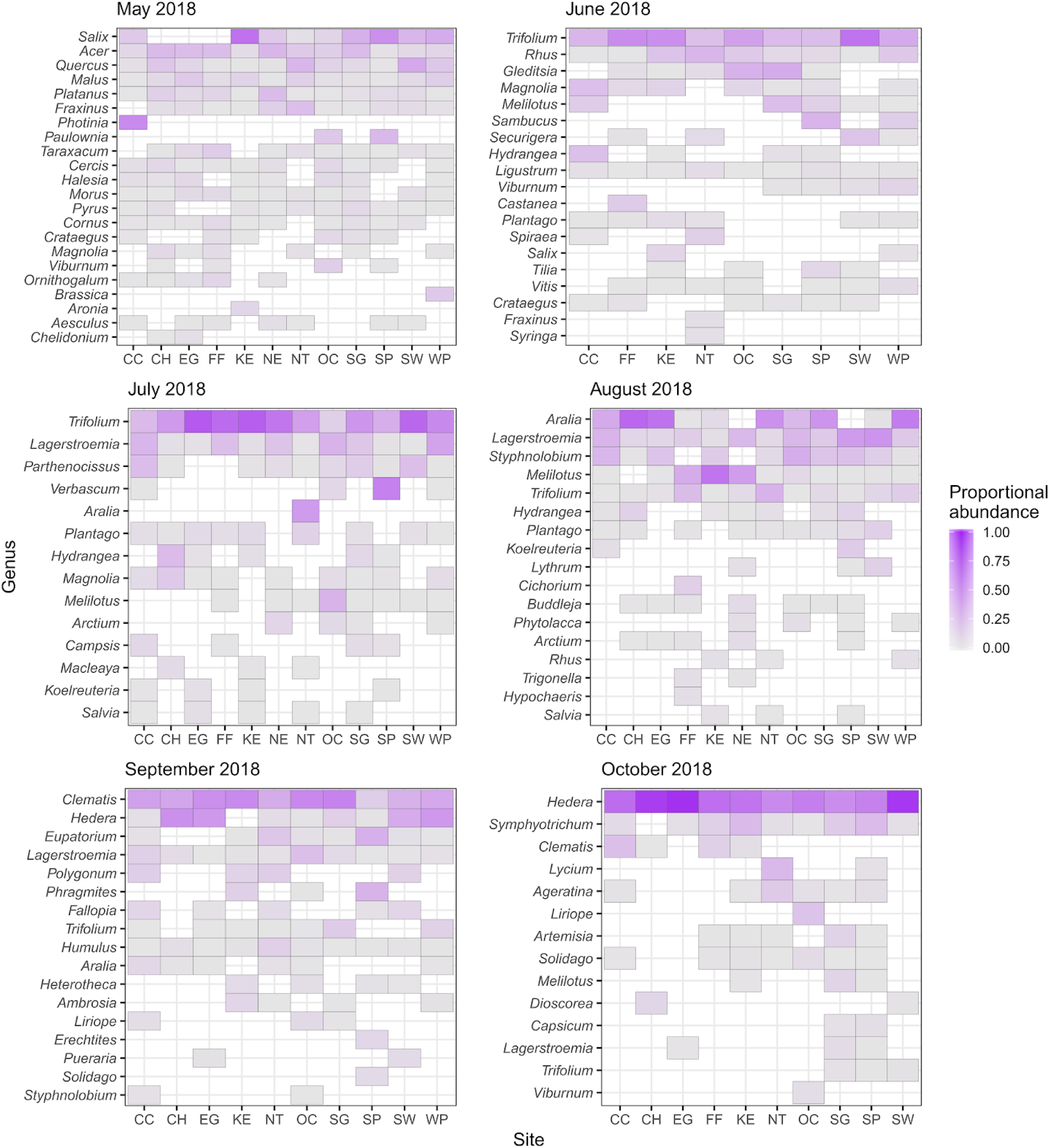
Genus composition of 2018 pollen samples. Genera are ranked by mean proportional abundance across sites. Only genera with a maximum within-site proportional abundance of at least 2.5% are shown.

In July **(Figures 6, 7)**, both weight gain and genus richness were attenuated **(Figures 3, 4)**, and pollen composition became dominated by clover (*Trifolium*) and Virginia creeper (*Parthenocissus*), accompanied by a strong presence of sweetclover (*Melilotus*) in 2017 and ornamental crepe myrtle (*Lagerstroemia*) in 2018. Mullein (*Verbascum*) and magnolia (*Magnolia*) were also important pollen sources, particularly in 2017.

During the apparently sparse month of August **(Figures 3, 6, 7)**, the abundance of clover (*Trifolium*) pollen declined but sweetclover (*Melilotus*) remained abundant. Spikenard (most likely the invasive understory tree, *Aralia elata*) emerged as a major pollen source. Crepe myrtle (*Lagerstroemia*) continued to be prominent alongside another ornamental species, Japanese pagoda tree (*Styphnolobium japonicum* is the only member of its genus present in our study area), which is commonly planted as a street tree in our study system and has become naturalized in some forested areas. The August-September interval saw another strong discontinuity in pollen composition **(Figure 5)**.

In September **(Figures 6, 7)**, clematis (most likely sweet autumn clematis, *Clematis ternifolia*) and ivy (*Hedera helix* and/or *H. hibernica*) dominated pollen samples, accompanied in 2017 by a strong presence of elm (the Chinese elm, *Ulmus parviflora*, is the only member of its genus that flowers in September in our study system). In 2017, crepe myrtle (*Lagerstroemia*) remained a major resource. Boneset (*Eupatorium*) was found across most sites in both years but was especially abundant in 2017.

At the close of the foraging season in October (sampled only in 2018) **(Figure 7)**, pollen samples from all sites consisted mainly of ivy (*H. helix* and/or *H. hibernica*). Late season asters were also common, though, including American asters (*Symphyotrichum*), snakeroot (*Ageratina*), *Artemisia* (likely *A. vulgaris*, common mugwort), and goldenrod (*Solidago*), and clematis remained important at some sites.

### Trait analysis

Trait-based patterns were generally consistent across the two years of our study **(Figure 8)**. With respect to growth habit, we saw a predominance of trees and shrubs early in the season, giving way to herbs in mid-summer, then returning to trees and shrubs in August, and finally shifting to vines in September and October. In terms of life cycle, the contribution of perennials appears to have dwarfed that of biennials and annuals throughout the whole foraging season, with the caveat that some sampling periods had a large proportion of genera with undetermined life cycle due to trait non-uniformity among congeners. Native genera ostensibly comprised the majority of pollen in May and June while exotic genera predominated in July, August, September, and October. These patterns should be interpreted cautiously, though, since the proportion of undetermined genera was high for most sampling periods, and in some cases abundant genera that could represent both native and exotic species (e.g. *Aralia, Clematis, Phragmites*) likely consisted mainly of their exotic constituents (e.g. *A. elata, C. ternifolia, P. australis*).

**Figure 8:**
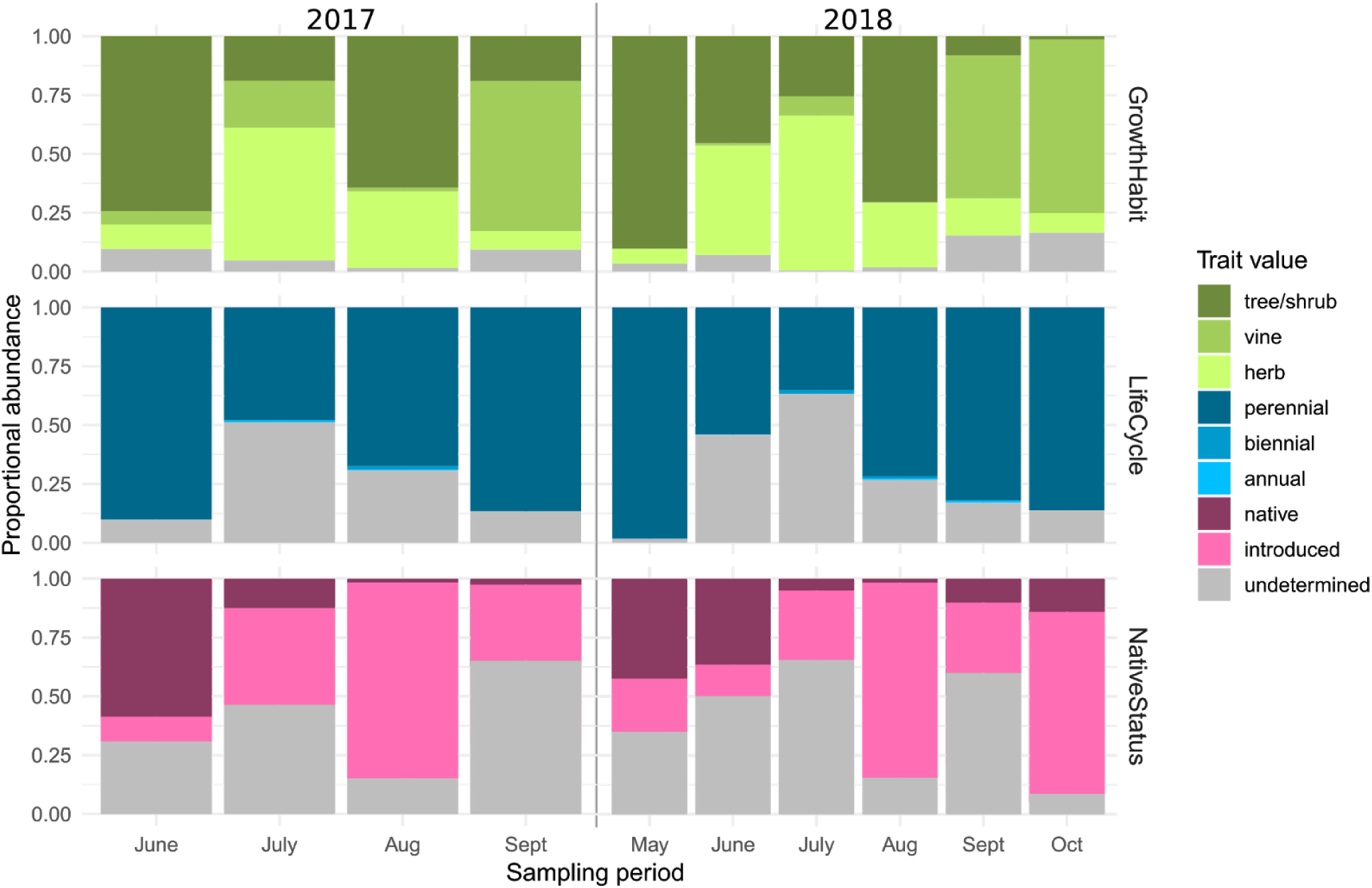
Trait composition by sampling period. Traits were scored as undetermined for a given genus when they were non-uniform across constituent species.

## Discussion

Continuous colony weight monitoring revealed strong gain and loss motifs. We interpret these dynamics as a measure of the overall abundance of floral resources within the colonies’ foraging range, with the caveat that colony weight is affected by overall health and honey bee life history in addition to resource availability. We mitigated these potentially confounding effects by omitting colonies with health anomalies, standardizing colony starting conditions, and analyzing detrended weight dynamics (i.e. differenced time series) rather than absolute weight. Under this interpretation, spring in our study system is a time of plenty, characterized by two pulses of resource availability, the first beginning in mid-May and the second in mid-June. Resource availability tapers off through July and reaches a minimum in August, a period during which all colonies in our study lost weight. After a brief pulse of resource availability in early September, the growing season is effectively over, with minimal resource availability from mid-September to the end of our measuring period in mid-October. These results--especially the occurrence of a late-summer/early-fall dearth of floral resources--are broadly consistent with temporal patterns of floral resource availability documented in England (Couvillon et al. 2014), France (Requier et al. 2015), Denmark (Lecocq et al. 2015), and Germany (Danner et al. 2017). That such similar results should be found in study systems on opposite sides of the Atlantic might be explained in part by the fact that several of the most important genera in our study system (e.g. *Trifolium, Melilotus, Hedera*) are European introductions that can also be major floral resources in their native range. The notion of a “summer dearth”, however, is a ubiquitous anecdote among beekeepers in temperate ecoregions [e.g. Japan: (Hiratsuka 1920), Ireland (Morony 1906), USA (Demuth 1918)], and the dynamics we observed may be broadly characteristic of temperate floral communities. This would suggest that urban landscapes, despite the extreme modification of their floral communities, conform to archetypal patterns of floral phenology that are shaped more by climate than by the particulars of individual landscapes.

One of the most salient patterns in our data is the importance of woody perennials, including both trees/shrubs and woody vines like *Hedera, Parthenocissus*, and *Clematis*. These findings contribute to a growing consensus about the importance of woody plants as pollinator resources (Mach and Potter 2018, Donkersley 2019), perhaps especially in urban environments (Burgett et al. 1978, Wattles 2017, Lau et al. 2019). In May, the most important resources included wind-pollinated trees like *Salix, Acer, Quercus, Fraxinus*, and *Platanus*, reinforcing Saunders’ (2018) call for increased attention to be given to wind-pollinated plants as resources for pollinators. Later in the season, ornamental trees and shrubs like *Lagerstroemia* and *Styphnolobium*, along with adventitious shrubs and woody vines like *Aralia, Parthenocissus, Clematis*, and *Hedera*, account for the majority of pollen foraging. Only in July did herbaceous plants consistently account for the majority of pollen foraging, and this was attributable mostly to a single dominant genus (*Trifolium*). These findings highlight the importance of accounting for trees and other woody plants in studies of plant-pollinator ecology, particularly in urban areas where these resources seem to be of prime importance, despite the technical challenges of canopy sampling that have prompted some authors to focus exclusively on low vegetation (Matteson et al. 2013, Lowenstein et al. 2019).

The strongest discontinuities in the composition of our pollen samples, particularly evident in clustering patterns seen in PCoA ordination **(Figure 5)**, occurred between the May and June sampling periods and between the August and September sampling periods. As observed by Robertson (1895), the late-spring discontinuity reflects the closure of the tree canopy, which marks the end of flowering for most tree and understory species, with a corresponding shift of pollinator visitation to edge and open habitats. The late-summer discontinuity in our data probably reflects the effective (though not absolute) end of *Trifolium* and *Melilotus* bloom (which may also account for the observed weight loss of colonies in August) and the shift to the fall flora dominated by *Clematis* and *Hedera*.

An unexpected--and perhaps characteristically urban--pattern in our data is the overwhelming abundance of woody vines in summer and fall pollen samples, beginning in July with *Parthenocissus* and peaking in September and October with *Clematis* and *Hedera*. In our study region, *Parthenocissus* is represented by both native *P. quinquefolia* and exotic *P. tricuspidata* (*USDA, NRCS 2019). The only species of Clematis* abundant in our study system that flowers in the fall is exotic *C. ternifolia* (D. Sponsler, personal observation). *Hedera* is represented by]*H. helix* and *H. hibernica*, both exotic (USDA, NRCS 2019). With the exception of the native *P. quinquefolia*, all these vines were introduced to North America as ornamentals and have since become widely naturalized in urban and suburban areas, where they thrive ubiquitously in both seminatural and developed areas. It is puzzling, in light of our findings, that neither *Clematis* nor *Hedera* is mentioned in classic surveys of North American pollinator-attractive flora, and *Parthenocissus* is deemed at best only a marginal nectar source of regional importance (Lovell 1918, Pellett 1920, Ayers and Harman 1992). This raises the question of whether the dominance of these vines as summer and fall floral resources was simply overlooked in the past--reflecting the general neglect of urban landscapes in 20th-century literature (Pickett et al. 2016)--or whether it is the result of more recent invasive success. In either case, if these taxa dominate flower-insect interactions in other cities to the extent observed here, it would be a striking case of urban biotic homogenization (McKinney 2006) propagated from the level of community composition to that of interaction network structure. Indeed, at least in the case of *Hedera*, its comparable importance as a bee plant in our study system and in Brighton, UK (Garbuzov and Ratnieks 2014), together with its widespread distribution in North American cities and suburbs (Jones and Reichard 2009), suggest that its role as a homogenizer of urban ecosystems may already be well-established. It should also be noted that *Clematis* and *Hedera* dominate the composition of our pollen samples at a time of year traditionally associated with the flowering of native asters (e.g. *Ageratina, Cichorium, Eupatorium, Eutrochium, Solidago, Symphyotrichum*) (Lovell 1918, Pellett 1920, Ayers and Harman 1992) and still dominated by native asters in rural areas (Sponsler et al. 2017); while detected in our samples, these native flora were dwarfed in their significance by *Clematis* and *Hedera*.

The special importance of the exotic vines in our data reflects a larger pattern in which exotic plants play a prominent role in the floral resource community of our study system. With the caveat that our genus trait analysis was hampered by trait non-uniformity in many genera, our data suggest a seasonal structure to the relative importance of native and exotic flora **(Figure 8)**. Native flora predominate early in the season, corresponding to the flowering of native trees and shrubs. In July and August, a season of overall dearth in floral resources **(Figure 3)**, exotic plants predominate. This pattern could be interpreted not as a displacement of native floral resources but a compensation for a period of natural scarcity. The ornamental trees *Lagerstroemia* and *Styphnolobium*, absent from previous surveys of North American floral resources (Lovell 1918, Pellett 1920, Ayers and Harman 1992), deserve special mention since they are not generally regarded as invasive. Moreover, the evident function of these ornamental trees as resources for flower-visiting insects suggests a caveat to Quigley’s (2011) dismissal of designed greenspaces as “Potempkin gardens” devoid of ecological function. While the annual ornamentals Quigley probably had in mind appear, indeed, to be of negligible importance as pollen resources **(Figure 8)**, their woody counterparts (e.g. *Lagerstoemia, Styphnolobium, Ligustrum, Hydrangea, Photinia, Magnolia, Phellodendron*) can be major pollen resources. As discussed above, however, the predominance of exotic flora--specifically *Clematis* and *Hedera*--continues through September and October, where their relative abundance could be interpreted as an ecological displacement of native asters. Notably, while annuals were virtually absent from our pollen samples, there is some evidence that annual ornamentals can be significant late-season nectar sources (Erickson et al. 2019).

The relationship between colony weight dynamics and pollen sample composition requires careful interpretation. First, any relationship between our weight data and pollen data is partly obscured by the fact that our weight data are continuous but our pollen data represent only a several-day interval per sampling period. This is a limitation that could theoretically be overcome by higher sampling frequency. A more fundamental constraint on the joint interpretation of weight and pollen data is that honey bee colony weight is driven mainly by changes in stored honey (i.e. the product of nectar foraging), but pollen trap samples represent only pollen foraging. For taxa thought to be major sources of both pollen and nectar (e.g. *Trifolium, Melilotus*) (Ayers and Harman 1992), relative abundance in pollen samples may approximate simultaneous contribution to nectar foraging. Other taxa, such as the wind-pollinated trees that dominate our May samples, produce copious pollen but little or no nectar (Ayers and Harman 1992). Conversely, taxa such as knotweed (*Reynoutria*) are prolific nectar sources but not thought to contribute significantly to pollen foraging (Ayers and Harman 1992). Thus, we recommend the conservative approach of interpreting pollen samples as qualitative indicators of flora in bloom during contemporaneous patterns of weight dynamics, with further interpretation based on ancillary data or expert knowledge of the study system.

While the focus of our study is on the use of honey bees as a model generalist forager, our data have practical relevance for beekeepers in study region. Colony weight dynamics can inform hive inspection regimes and allow beekeepers to plan management interventions such as “supering” (the addition of boxes to accomodate colony growth), honey harvest, supplemental feeding, or colony splitting (Gary 1992a). This of particular importance to urban beekeepers who often contend with difficult access to their apiaries (Alton and Ratnieks 2016) and must schedule their management judiciously. It is also important to note times of dearth so that inter-colony robbing can be mitigated, e.g. through the use of entrance reducers and the minimizing of open-hive manipulations (Gary 1992b). The clear demarcation of spring, summer, and fall floral communities in our data is also of apicultural significance. Traditionally, apiarists in our study region recognize spring and fall nectar flows, and honey is often marketed in terms of these seasonal varieties. Our data show a more nuanced, three-season pattern of nectar flows. Knowing this pattern would potentially allow beekeepers to harvest a third seasonal honey variety, a significant asset to urban beekeepers who lack access to single-species cropping needed to market floral varietal honeys.

### Conclusions and future research

Frankl et al. (2005) pose the question of whether the landscape-scale quantification of floral resources is possible. While there may be multiple ways to answer this question in the affirmative, including the approach presented by its original authors (Frankl et al. 2005), we submit that our “honey bee foraging assay”--the combination of continuous weight monitoring and pollen metabarcoding--constitutes a landscape-scale quantification of floral resources. We acknowledge that honey bee foraging is not an unbiased sampling of floral resources, and we do not interpret our results as an objective census of the floral community. Nevertheless, an objective census of floral resources would arguably be less relevant to questions of ecological function than the “biased” sampling of a generalist florivore. In principle, the foraging assay approach to landscape-scale floral resource surveying could be implemented with model organisms other than the honey bee, and we urge the development of such methods. Bumble bees (*Bombus* spp.), in particular, are amenable to both pollen barcoding (Pornon et al. 2016) and colony weight monitoring (Goulson et al. 2002), and their choice of floral resources would be expected to differ from that of honey bees, though with substantial overlap (Leonhardt and Blüthgen 2012).

It is not incidental that our methodology was implemented in a complex urban landscape. With respect to the ecology and conservation of plant-pollinator communities, the question is no longer *whether* cities can support such communities (Baldock et al. 2015) but rather *how* they are structured by the heterogeneous habitats of which cities are composed (Baldock et al. 2019). Our work begins to shed light on this question, revealing the emergence of weedy vines as dominant floral resources, the seasonal structuring of the relative importance of trait-based floral guilds (e.g. native vs. exotic, woody vs. herbaceous), and the dramatic temporal dynamics of overall floral resource availability. While we urge caution in extrapolating our findings beyond our Philadelphia study system, the general consistency of our findings with comparable datasets (Garbuzov and Ratnieks 2014, Couvillon et al. 2014, Lecocq et al. 2015, Requier et al. 2015, Danner et al. 2017, Lau et al. 2019) suggests that the patterns seen in our data are shaped by ecological and social drivers that are not unique to our locality. Future work should focus on elucidating these ecological drivers and formulating testable hypotheses regarding the relationships between landscape, floral resources, and florivores.

## Acknowledgements

T. Jones, D. Brough, K. Oxman assisted and field and laboratory work. K. Frank, K. Wattles and members of the Philadelphia Beekeepers Guild and the Philadelphia Botanical Club provided informative conversation regarding local floral resources. M. McIntyre, Weavers Way Coop, Congregation Rodeph Shalom, Hotel Sofitel, Shane Confectionery, Greensgrow Farm, The Philadelphia Insectarium, the Philadelphia Business and Technology Center, Mt. Moriah Cemetery, Share Food Program, W. B. Saul high school, Paradiso Restaurant, and A. Pfeffer provided apiary locations for this study. This work was funded by a USDA-NIFA postdoctoral fellowship for D. Sponsler (grant 2017-07141), an Apes Valentes research award to D. Sponsler, and USDA–NIFA–SCRI grant 2016–51181–235399, “Protecting Pollinators with Economically Feasible and Environmentally Sound Ornamental Horticulture.”

## References

Alton, K., and F. Ratnieks. 2016. Roof Top Hives: Practical Beekeeping or Publicity Stunt? Bee World 93:64–67.

Aronson, M. F. J., F. A. La Sorte, C. H. Nilon, M. Katti, M. A. Goddard, C. A. Lepczyk, P. S. Warren, N. S. G. Williams, S. Cilliers, B. Clarkson, C. Dobbs, R. Dolan, M. Hedblom, S. Klotz, J. L. Kooijmans, I. Kühn, I. Macgregor-Fors, M. McDonnell, U. Mörtberg, P. Pysek, S. Siebert, J. Sushinsky, P. Werner, and M. Winter. 2014. A global analysis of the impacts of urbanization on bird and plant diversity reveals key anthropogenic drivers. Proceedings of the Royal Society B 281:20133330.

Ayers, G. S., and J. R. Harman. 1992. Bee forage of North America and the potential for planting for bees. Pages 437–493 in J. M. Graham, editor. The Hive and the Honey Bee. Dadant & Sons, Hamilton, IL.

Baldock, K. C. R., M. A. Goddard, D. M. Hicks, W. E. Kunin, N. Mitschunas, H. Morse, L. M. Osgathorpe, S. G. Potts, K. M. Robertson, A. V. Scott, P. P. A. Staniczenko, G. N. Stone, I. P. Vaughan, and J. Memmott. 2019. A systems approach reveals urban pollinator hotspots and conservation opportunities. Nature Ecology & Evolution 3:363–373.

Baldock, K. C. R., M. A. Goddard, D. M. Hicks, W. E. Kunin, N. Mitschunas, L. M. Osgathorpe, S. G. Potts, K. M. Robertson, A. V. Scott, G. N. Stone, I. P. Vaughan, and J. Memmott. 2015. Where is the UK’s pollinator biodiversity? The importance of urban areas for flower-visiting insects. Proceedings of the Royal Society B 282:20142849.

Berry, D., K. Ben Mahfoudh, M. Wagner, and A. Loy. 2011. Barcoded Primers Used in Multiplex Amplicon Pyrosequencing Bias Amplification. Applied and Environmental Microbiology 77:7846–7849.

Burgett M, Caron DM, Ambrose JT. 1978. Urban apiculture. In: Frankie GW, Kohler CS, eds. Perspectives in Urban Entomology. New York: Academic Press, 187–219.

Cheng, T., C. Xu, L. Lei, C. Li, Y. Zhang, and S. Zhou. 2016. Barcoding the kingdom Plantae: new PCR primers for ITS regions of plants with improved universality and specificity. Molecular Ecology Resources 16:138–149.

Chen, S., H. Yao, J. Han, C. Liu, J. Song, L. Shi, Y. Zhu, X. Ma, T. Gao, X. Pang, K. Luo, Y. Li, X. Li, X. Jia, Y. Lin, and C. Leon. 2010. Validation of the ITS2 Region as a Novel DNA Barcode for Identifying Medicinal Plant Species. PloS One 5:e8613.

Chesapeake Bay Conservancy. 2017. Land cover data project.

Clemants, S., and G. Moore. 2003. Patterns of Species Richness in Eight Northeastern United States Cities. Urban Habitats 1:4–16.

Couvillon, M. J., R. Schürch, and F. L. W. Ratnieks. 2014. Waggle Dance Distances as Integrative Indicators of Seasonal Foraging Challenges. PloS One 9:e93495.

Danner, N., A. Keller, S. Härtel, and I. Steffan-Dewenter. 2017. Honey bee foraging ecology: Season but not landscape diversity shapes the amount and diversity of collected pollen. PloS One 12:e0183716.

Demuth, B. M. 1918. Bees for the winter cluster. Gleanings in Bee Culture 46:462–463.

Donkersley, P. 2019. Trees for bees. Agriculture, ecosystems & environment 270-271:79–83.

Erickson, E., Adam. S., Russo, L., Wojcik, V., Patch, H.M., and C.M. Grozinger. 2019. More than meets the eye: The role of ornamental plants in supporting pollinators. Environmental Entomology (in press).

Esling, P., F. Lejzerowicz, and J. Pawlowski. 2015. Accurate multiplexing and filtering for high-throughput amplicon-sequencing. Nucleic Acids Research 43:2513–2524.

Fenneman, F., and D. M. Johnson. 1946. Physiographic divisions of the conterminous U. S. United States Geological Survey.

Frankl, R., S. Wanning, and R. Braun. 2005. Quantitative floral phenology at the landscape scale: Is a comparative spatio-temporal description of “flowering landscapes” possible? Journal for Nature Conservation 13:219–229.

Garbuzov, M., and F. L. W. Ratnieks. 2014. Ivy: an underappreciated key resource to flower-visiting insects in autumn. Insect Conservation and Diversity 7:91–102.

Gary, N. E. 1992a. Activities and behavior of honey bees. Pages 269–372 in J. M. Graham, editor. The Hive and the Honey Bee. Dadant & Sons, Hamilton, IL.

Gary, N. E. 1992b. Activities and behavior of honey bees. Pages 269–372 in J. M. Graham, editor. The Hive and the Honey Bee. Dadant & Sons, Hamilton, IL.

Geslin, B., B. Gauzens, E. Thébault, and I. Dajoz. 2013. Plant Pollinator Networks along a Gradient of Urbanisation. PloS one 8:e63421.

Gordon, A., G. J. Hannon, and Others. 2010. Fastx-toolkit. FASTQ/A short-reads preprocessing tools. http://hannonlab.cshl.edu/fastx_toolkit5.

Goulson, D. 2003. Effects of Introduced Bees on Native Ecosystems. Annual Review of Ecology, Evolution, and Systematics 34:1–26.

Goulson, D., W. Hughes, L. Derwent, and J. Stout. 2002. Colony growth of the bumblebee, Bombus terrestris, in improved and conventional agricultural and suburban habitats. Oecologia 130:267–273.

Hiratsuka, Y. 1920. Honey plants in Japan. American Bee Journal 60:100.

Jones, C. C., and S. Reichard. 2009. Current and Potential Distributions of Three Non-Native Invasive Plants in the Contiguous USA. Natural Areas Journal 29:332–343.

Keller, A., N. Danner, G. Grimmer, V. der Ankenbrand M., V. der Ohe K., W. Ohe, S. Rost, S. Härtel, and I. Steffan-Dewenter. 2015. Evaluating multiplexed next-generation sequencing as a method in palynology for mixed pollen samples. Plant Biology 17:558–566.

Kozich, J. J., S. L. Westcott, N. T. Baxter, S. K. Highlander, and P. D. Schloss. 2013. Development of a dual-index sequencing strategy and curation pipeline for analyzing amplicon sequence data on the MiSeq Illumina sequencing platform. Applied and Environmental Microbiology 79:5112–5120.

Kraaijeveld, K., L. A. Weger, M. Ventayol García, H. Buermans, J. Frank, P. S. Hiemstra, and J. T. Dunnen. 2015. Efficient and sensitive identification and quantification of airborne pollen using next-generation DNA sequencing. Molecular Ecology Resources 15:8–16.

Lau, P., V. Bryant, J. D. Ellis, Z. Y. Huang, J. Sullivan, D. R. Schmehl, A. R. Cabrera, and J. Rangel. 2019. Seasonal variation of pollen collected by honey bees (Apis mellifera) in developed areas across four regions in the United States. PloS One 14:e0217294.

Lecocq, A., P. Kryger, F. Vejsnæs, and A. Bruun Jensen. 2015. Weight Watching and the Effect of Landscape on Honeybee Colony Productivity: Investigating the Value of Colony Weight Monitoring for the Beekeeping Industry. PloS One 10:e0132473.

Legendre, P., and E. D. Gallagher. 2001. Ecologically meaningful transformations for ordination of species data. Oecologia 129:271–280.

Leonhardt, S. D., and N. Blüthgen. 2012. The same, but different: pollen foraging in honeybee and bumblebee colonies. Apidologie 43:449–464.

Lovell, J. H. 1918. The Flower and the Bee: Plant Life and Pollination. C. Scribner’s sons.

Lowenstein, D. M., K. C. Matteson, and E. S. Minor. 2019. Evaluating the dependence of urban pollinators on ornamental, non-native, and “weedy” floral resources. Urban Ecosystems 22:293–302.

Mach, B. M., and D. A. Potter. 2018. Quantifying bee assemblages and attractiveness of flowering woody landscape plants for urban pollinator conservation. PloS One 13:e0208428.

Matteson, K. C., J. B. Grace, and E. S. Minor. 2013. Direct and indirect effects of land use on floral resources and flower-visiting insects across an urban landscape. Oikos 122:682–694.

McKinney, M. L. 2006/1. Urbanization as a major cause of biotic homogenization. Biological Conservation 127:247–260.

Morony, W. 1906. The month’s work. Irish Bee Journal 5:35.

USDA, NRCS. 2019. The PLANTS Database (http://plants.usda.gov). National Plant Data Team, Greensboro, NC 27401-4901 USA.

Oksanen, J., G. Blanchet, M. Friendly, R. Kindt, P. Legendre, D. McGlinn, and Others. 2019. vegan: Community ecology package. R package version 2.5-6. 2019.

Pedersen, E. J., D. L. Miller, G. L. Simpson, and N. Ross. 2019. Hierarchical generalized additive models in ecology: an introduction with mgcv. PeerJ 7:e6876.

Pellett, F. C. 1920. American honey plants; together with those which are of special value to the beekeeper as sources of pollen. Hamilton: Dadant and Sons. Available at https://archive.org/details/americanhoney00pell

Pickett, S. T. A., M. L. Cadenasso, D. L. Childers, M. J. Mcdonnell, and W. Zhou. 2016. Evolution and future of urban ecological science: ecology in, of, and for the city. Ecosystem Health and Sustainability 2:e01229.

Pornon, A., N. Escaravage, M. Burrus, H. Holota, A. Khimoun, J. Mariette, C. Pellizzari, A. Iribar, R. Etienne, P. Taberlet, M. Vidal, P. Winterton, L. Zinger, and C. Andalo. 2016. Using metabarcoding to reveal and quantify plant-pollinator interactions. Scientific Reports 6:27282.

Quigley, M. F. 2011. Potemkin gardens: Biodiversity in small designed landscapes. Pages 85–91 in J. Niemelä, editor. Urban Ecology: Patterns, Processes, and Applications. Oxford University Press.

R Core Team. 2019.

Requier, F., J.-F. Odoux, T. Tamic, N. Moreau, M. Henry, A. Decourtye, and V. Bretagnolle. 2015. Honey bee diet in intensive farmland habitats reveals an unexpectedly high flower richness and a major role of weeds. Ecological Applications 25:881–890.

Richardson, R. T., H. R. Curtis, E. G. Matcham, C.-H. Lin, S. Suresh, D. B. Sponsler, L. E. Hearon, and R. M. Johnson. 2019a. Quantitative multi-locus metabarcoding and waggle dance interpretation reveal honey bee spring foraging patterns in Midwest agroecosystems. Molecular Ecology 28:686–697.

Richardson, R. T., C.-H. Lin, J. O. Quijia, N. S. Riusech, K. Goodell, and R. M. Johnson. 2015. Rank-based characterization of pollen assemblages collected by honey bees using a multi-locus metabarcoding approach. Applications in Plant Sciences 3:1500043.

Richardson, R. T., D. B. Sponsler, H. McMinn-Sauder, and R. M. Johnson. 2019b. MetaCurator: A hidden Markov model-based toolkit for extracting and curating sequences from taxonomically-informative genetic markers. Methods in Ecology and Evolution 2019;00:1–6.

Robertson, C. 1895. The Philosophy of Flower Seasons, and the Phaenological Relations of the Entomophilous Flora and the Anthophilous Insect Fauna. The American Naturalist 29:97–117.

Rognes, T., T. Flouri, B. Nichols, C. Quince, and F. Mahé. 2016. VSEARCH: a versatile open source tool for metagenomics. PeerJ 4:e2584.

Saunders, M. E. 2018. Insect pollinators collect pollen from wind-pollinated plants: implications for pollination ecology and sustainable agriculture. Insect Conservation and Diversity 11:13–31.

Schnell, I. B., K. Bohmann, and M. T. P. Gilbert. 2015. Tag jumps illuminated--reducing sequence-to-sample misidentifications in metabarcoding studies. Molecular Ecology Resources 15:1289–1303.

Sickel, W., M. J. Ankenbrand, G. Grimmer, A. Holzschuh, S. Härtel, J. Lanzen, I. Steffan-Dewenter, and A. Keller. 2015. Increased efficiency in identifying mixed pollen samples by meta-barcoding with a dual-indexing approach. BMC Ecology 15:20.

Smart, M. D., R. S. Cornman, D. D. Iwanowicz, M. McDermott-Kubeczko, J. S. Pettis, M. S. Spivak, and C. R. V. Otto. 2017. A Comparison of Honey Bee-Collected Pollen From Working Agricultural Lands Using Light Microscopy and ITS Metabarcoding. Environmental Entomology 2017:1–12.

Sponsler, D. B., E. G. Matcham, C.-H. Lin, J. L. Lanterman, and R. M. Johnson. 2017. Spatial and taxonomic patterns of honey bee foraging: A choice test between urban and agricultural landscapes. Journal of Urban Ecology 2017:1–7.

Taberlet, P., E. Coissac, F. Pompanon, L. Gielly, C. Miquel, A. Valentini, T. Vermat, G. Corthier, C. Brochmann, and E. Willerslev. 2007. Power and limitations of the chloroplast trnL (UAA) intron for plant DNA barcoding. Nucleic Acids Research 35:e14.

Visscher, P. K., and T. D. Seeley. 1982. Foraging Strategy of Honeybee Colonies in a Temperate Deciduous Forest. Ecology 63:1790–1801.

Wang, X.-C., C. Liu, L. Huang, J. Bengtsson-Palme, H. Chen, J.-H. Zhang, D. Cai, and J.-Q. Li. 2015. ITS1: a DNA barcode better than ITS2 in eukaryotes? Molecular Ecology Resources 15:573–586.

Wardhaugh, C. W. 2015. How many species of arthropods visit flowers? Arthropod-Plant Interactions 9:547–565.

Wattles, K. 2017. Trees and Shrubs for Bees. Available at: https://phillybeekeepers.org/wp-content/uploads/2017/02/BToS_3rd-ed_20170204_5pp.pdf

Wickham, H. 2017. Package tidyverse: Easily install and load the “Tidyverse.” https://cran.r-project.org/web/packages/tidyverse/index.html

Wickham, H., R. Francois, L. Henry, K. Müller, and Others. 2015. dplyr: A grammar of data manipulation. R package version 0. 4 3.

Wood, S. N. 2017. Generalized Additive Models: An Introduction with R, Second Edition. Chapman and Hall/CRC.

Zhang, J., K. Kobert, T. Flouri, and A. Stamatakis. 2014. PEAR: a fast and accurate Illumina Paired-End reAd mergeR. Bioinformatics 30:614–620.

